# Parsnp 2.0: Scalable Core-Genome Alignment for Massive Microbial Datasets

**DOI:** 10.1101/2024.01.30.577458

**Authors:** Bryce Kille, Michael G. Nute, Victor Huang, Eddie Kim, Adam M. Phillippy, Todd J. Treangen

## Abstract

**Motivation:** Since 2016, the number of microbial species with available reference genomes in NCBI has more than tripled. Multiple genome alignment, the process of identifying nucleotides across multiple genomes which share a common ancestor, is used as the input to numerous downstream comparative analysis methods. Parsnp is one of the few multiple genome alignment methods able to scale to the current era of genomic data; however, there has been no major release since its initial release in 2014.

**Results:** To address this gap, we developed Parsnp v2, which significantly improves on its original release. Parsnp v2 provides users with more control over executions of the program, allowing Parsnp to be better tailored for different use-cases. We introduce a partitioning option to Parsnp, which allows the input to be broken up into multiple parallel alignment processes which are then combined into a final alignment. The partitioning option can reduce memory usage by over 4x and reduce runtime by over 2x, all while maintaining a precise core-genome alignment. The partitioning workflow is also less susceptible to complications caused by assembly artifacts and minor variation, as alignment anchors only need to be conserved within their partition and not across the entire input set. We highlight the performance on datasets involving thousands of bacterial and viral genomes.

**Availability:** Parsnp is available at https://github.com/marbl/parsnp

## 1 Introduction

As a result of the democratization of sequencing technology, the quality and quantity of publicly available assembled genomes has grown at an unprecedented rate [1, 2]. Multiple genome alignment, the process of identifying nucleotides which share a common ancestor across a set genomes, is a critical step in typical comparative genomics workflows. However, tools that aim to compute multiple genome alignments have struggled to keep up with the ever-increasing rate of growth of available data.

To reduce the complexity and maximize the signal of comparative genomics analysis for a group of closely-related genomes, it is often helpful to reduce them to their highly-conserved orthologous regions. By nature of being highly conserved and vertically inherited, orthologous regions are well suited for constructing phylogenies as well as identifying core functions and regulatory regions within microbial genomes.

While core-genome analysis has been enabled by many tools over the last decade, Parsnp [3] is one of the few capable of identifying and reporting the core genome and all of its variation from the sequences alone. Alternative methods, such as Roary [4] and Read2Tree [5], rely on genome annotation and annotated databases to map to orthologs and identify core-genome SNPs. Importantly, Parsnp constructs the core genome through multiple sequence alignment as opposed to pairwise alignment, therefore avoiding any reference bias.

To identify the core-regions of a set of genomes, Parsnp first identifies maximal unique matches (MUMs), sequences which are identical across all genomes and unique within each genome, to anchor the global core-genome alignment. Parsnp then identifies syntenic regions (locally collinear blocks) based on these MUMs, performs a recursive MUM search, and then passes each syntenic region on to multiple sequence alignment to obtain the SNPs for each genome.

Parsnp has been widely adopted since its release in 2014. Despite this, there have been no major releases for nearly a decade, leading to a consistent growth in open issues and feature-requests ranging from easier installation to new post-processing options. In addition, the recent influx of massive intra-species assemblies has identified a core issue in the Parsnp method: the length and quantity of MUMs often decreases as the number of input sequences grows due to minor variation and assembly artifacts. As a result, the identified core-regions in massive datasets, particularly those of low-quality or high diversity, are more fragmented and less abundant.

In a new release of Parsnp, we address a significant number of open issues and feature requests, fix subtle but impactful bugs, and provide a new divide-and-conquer workflow which allows Parsnp to identify core-regions which were not captured due to the aforementioned MUM dropout on massive datasets. In addition, our divide-and-conquer approach is embarrassingly parallel and requires substantially less memory than the original Parsnp workflow.

## 2 Methods

### 2.1 Partitioning workflow

As a core region in Parsnp is defined as a region that is present in all input sequences, any core region is also present in all sequences for any subset of the input. Parsnp v2 offers a partitioning workflow which takes advantage of this by partitioning the input genomes up into groups of equal size and running Parsnp v2 on each group, using the same reference for each partition. Then, Parsnp v2 reports the common regions across all partitioned core genomes as the final core genome.

The advantage of this workflow is that it is much more parallel and requires significantly less memory. In addition, it can lead to larger core genomes for very large input datasets since very large datasets typically suffer from MUM dropout, as mentioned previously.

### 2.1.1 Identifying final core-genome coordinates from partitions

In order to identify the coordinates of the final core genome with respect to a reference sequence, we compute the intersection of the aligned reference regions for each partition. Let *q* = *n/p* be the number of partitions of *n* query genomes with *p* genomes per partition and *R*_*i*_ = *{*(*b*_1_, *e*_1_), (*b*_2_, *e*_2_), …*}* be the begin and end coordinates of the reference for each of the LCBs in partition *i*.

We compute the interval intersection of all *q* interval sets to obtain the coordinates of the final core-genome intervals, *R*^*∗*^. The output alignments of each partition are then trimmed to match the coordinates of *R*^*∗*^. At the end of this step, each partition will have an alignment file with |*R*^*∗*^| LCBs, where each LCB corresponds to one of the intervals in *R*^*∗*^.

The trimmed output alignments of each partition are then stitched into a combined alignment of all input sequences, using the fact that the alignment columns represent an equivalence relationship between the reference and query sequences for each position. Special care must be taken for insertions with respect to the reference, though. To handle this particular case, we align inserted sequences across partitions using SPOA [6]. More details on this step can be found in the Supplementary.

### 2.2 Additional Parsnp v2 features

In addition to the aforementioned partitioning workflow and bug-fixes, there are many new improvements which greatly improve Parsnp’s utility. Until 2020, Parsnp was not available on the Bioconda channel and required a manual build of a custom MUSCLE library [7] in addition to the Parsnp binary. This, with the additional requirement of Python 2 and other dependencies not listed on Bioconda, made for a laborious and non-trivial installation process. In order to ameliorate these issues, we updated Parsnp to support Python 3, added Parsnp as well as its dependencies to Bioconda, and in effect rid the need for users to obtain and build Parsnp and its multiple dependencies.

We have also added numerous additional options to Parsnp’s interface which dictate the input, output, and internal pipeline of the Parsnp method. Users can now provide their query inputs through regular expression CLI matching and text files, output VCF files directly, use FastANI [8] for genome recruitment, use FastTree2 [9] for phylogeny reconstruction or skip the step as a whole, and more. These improvements greatly increase the utility of Parsnp v2, making it suitable for more custom end-user analysis.

## 3 Results

### 3.1 Performance and core-genome characteristics

All available complete assemblies for *S. aureus, M. tuberculosis, K. pneumoniae* were downloaded from NCBI. This resulted in groups of 2729, 828, and 4326 assemblies, respectively. In addition, we obtained a random sample of 100,000 complete SARS-CoV-2 genomes from GenBank, each containing at most 10 ambiguous nucleotides. For each group of assemblies, Parsnp (commit 1d3fbcc) was run on a server with 128GB of memory and 8 processors with partition sizes *p* of 50, 100, and 250 genomes as well as without partitioning (*p* = *n*). CoreDetector [11], a recent core-genome alignment method, was also run for comparison. As Parsnp has the convenience of producing .ggr files which can be viewed with Gingr, we demonstrate a visualization of the *M. tuberculosis* core genome in Figure 1.

**Figure 1:**
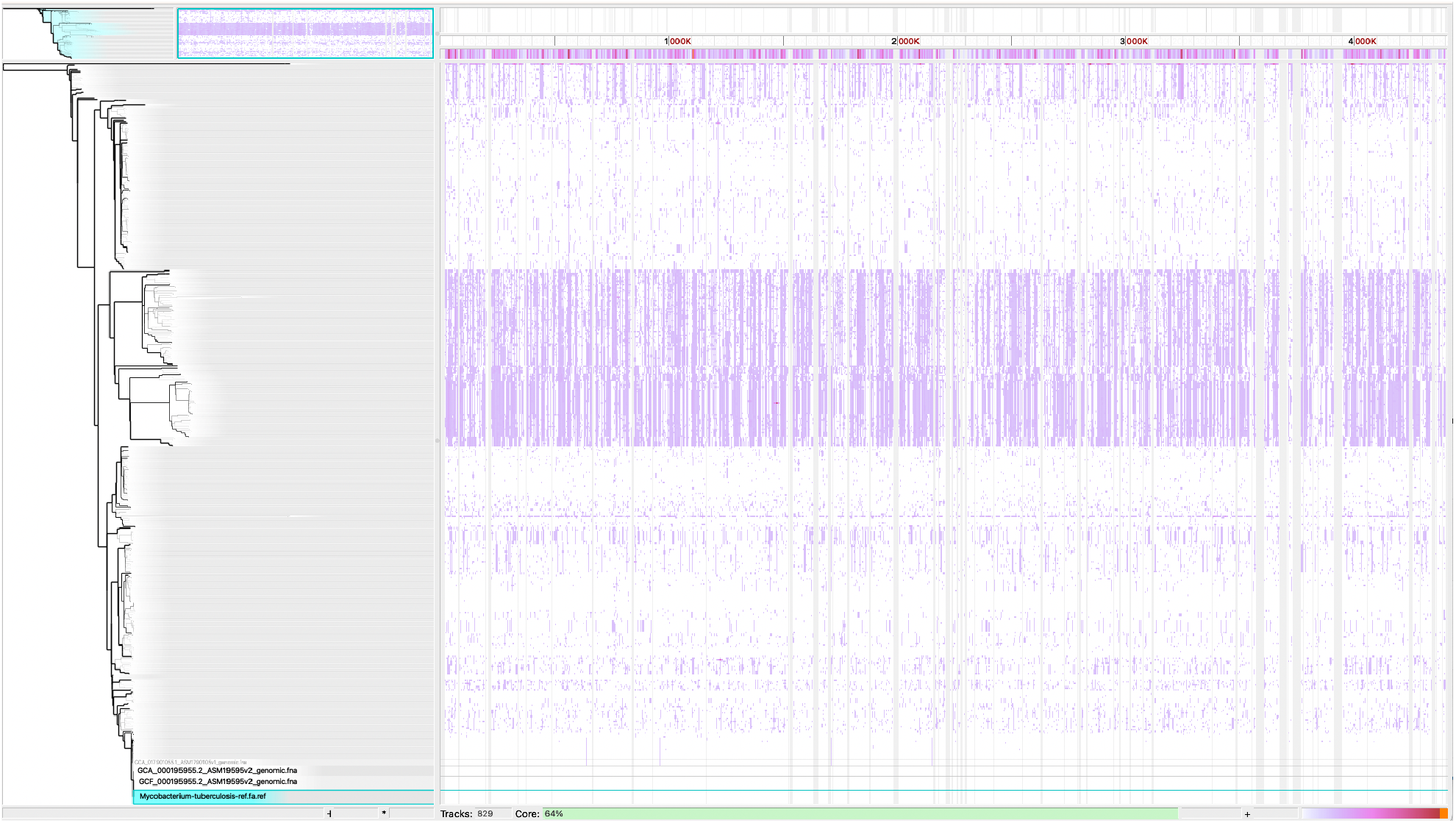
Gingr visualization of a Parsnp (*p* = 250) alignment on 828 *M. tuberculosis* genomes. When zoomed out, the visualization displays a heatmap of single-nucleotide variants. Base-level resolution views of the alignments become visible when inspecting smaller regions.

Large indels represent sequences not present in all of the input genomes and therefore are not considered part of the core genome, and as such, Parsnp will not report them. To enable a more appropriate comparison, LCBs with indels greater than 300 nucleotides were removed from CoreDetector’s output. After this adjustment, the size of the core region reported by CoreDetector was comparable to Parsnp, but the average divergence was consistently higher i.e. Parsnp was able to produce a better-aligned core genome of roughly the same size.

For each run, the peak memory usage as well as wall-clock time are reported in Table 1. In addition to the runtime performance, Table 1 also reports the weighted average divergence from the reference across all LCBs, where each LCB is weighted by the number of nucleotides it contains, the size of the core genome, and the number of LCBs.

**Table 1:** Performance of Parsnp on pangenomes of different sizes compared to CoreDetector. Parsnp was executed with different partition sizes *p* on each set of genomes. Total number of genomes *n* and dataset size shown is also shown. Mean reference divergence is defined as 1 minus the weighted average of the average nucleotide identity across all LCBs, where the weight is the total number of nucleotides in the LCB. The total core length is the number of reference nucleotides present in the core alignment, and the LCB count is the number of LCBs reported. The results of CoreDetector before removing LCBs with indels larger than 300bp are reported in parentheses (Parsnp did not produce any such LCBs). Both the non-partitioned run of Parsnp and CoreDetector failed on the *K. pneumoniae* due to excessive memory use. For SARS-CoV-2, the non-partitioned run of Parsnp failed to produce a core genome after 88 minutes due to lack of MUMs in the anchoring step and CoreDetector halted after 13 hours due to a runtime error.

| Dataset | Method | Wall Time (m) | CPU Time (m) | Memory (Gb) | Mean Reference % Divergence | Total Core Length (kbp) | LCB Count |
| --- | --- | --- | --- | --- | --- | --- | --- |
| *M. tuberculosis*<br>$n = 828$<br>3.5Gbp | Parsnp ($p = 50$) | 31 | 214 | **21.0** | 0.03 | 1727 | 4876 |
| | Parsnp ($p = 100$) | 38 | 253 | 46.3 | 0.03 | 2368 | 5995 |
| | Parsnp ($p = 250$) | 66 | 297 | 42.3 | 0.03 | 2776 | 4976 |
| | Parsnp ($p = n$) | 64 | 263 | 46.0 | 0.03 | 2997 | 4873 |
|  | CoreDetector | **25** | **64.7** | 25.5 | 0.07 (0.21) | 2669 (3231) | 1212 (1323) |
| *S. aureus*<br>$n = 2729$<br>7.4Gbp | Parsnp ($p = 50$) | **69** | 488 | **13.9** | 0.47 | 830 | 2725 |
| | Parsnp ($p = 100$) | 78 | 521 | 23.9 | 0.46 | 785 | 2421 |
| | Parsnp ($p = 250$) | 114 | 626 | 68.5 | 0.43 | 694 | 2064 |
| | Parsnp ($p = n$) | 188 | 857 | 96.9 | 0.38 | 414 | 1409 |
|  | CoreDetector | **69** | **163** | 54.2 | 0.50 (0.54) | 656 (898) | 457 (513) |
| *K. pneumoniae*<br>$n = 4326$<br>24Gbp | Parsnp ($p = 50$) | **183** | **1350** | **31.7** | 0.28 | 1314 | 6207 |
| | Parsnp ($p = 100$) | 203 | 1433 | 58.9 | 0.27 | 1258 | 5574 |
| | Parsnp ($p = 250$) | 304 | 1751 | 123.8 | 0.26 | 1138 | 4674 |
| | Parsnp ($p = n$) | NA | NA | > 128 | NA | NA | NA |
|  | CoreDetector | NA | NA | > 128 | NA | NA | NA |
| SARS-CoV-2<br>$n = 100000$<br>3.2Gbp | Parsnp ($p = 50$) | 55 | **313** | 7.76 | 0.15 | 25.7 | 134 |
| | Parsnp ($p = 100$) | **54** | 316 | 8.57 | 0.15 | 25.4 | 133 |
| | Parsnp ($p = 250$) | 61 | 380 | **6.89** | 0.15 | 25.0 | 123 |
| | Parsnp ($p = n$) | NA | NA | NA | NA | NA | NA |
|  | CoreDetector | NA | NA | NA | NA | NA | NA |

Table 1 shows that the memory and runtime advantages of the partitioned approach can be substantial and can enable alignment on hardware that might otherwise have been resource-prohibitive. While the wall time used by Parsnp (*p* = 50) and CoreDetector were comparable in the two datasets where CoreDetector completed successfully, CoreDetector required up to 3x less CPU time than Parsnp (*p* = 50). However, as most modern machines have at least 8 cores, the wall time is more representative of practical settings. For the partitioned mode of Parsnp, the memory usage scales linearly with both the partition size and the number of threads used; users can increase the number of threads used in return for shorter wall times so long as they have sufficient memory available.

In three of the four data sets, the partitioning results in a larger core genome with very little cost to the alignment quality. The remaining alignment for *M. tuberculosis* suggests that there are genomic datasets where partitioning decreases the core-genome coverage. One distinguishing feature of this dataset is that it contained more LCBs than the other datasets (Table 1) despite having fewer genomes and lower divergence. This has been observed in a prior study on *M. tuberculosis* genome alignments [12] as well.

Non-partitioned Parsnp (*p* = *n*) is likely at an advantage in this scenario; higher LCB counts due to high levels of rearrangements (and other factors) result in an increase in the number of synteny breakpoints. Non-partitioned Parsnp has a global view across all input genomes and is able to both filter out potential spurious MUMs and perform a recursive MUM search across all genomes to help mitigate the effect of high LCB counts. Conversely, the partitioned version is only aware of the LCB breakpoints within a given partition, so when merging the intervals across all partitions with high LCB counts, the core genome can degrade in size (Table 1). Better understanding of this phenomenon will require additional experiments on a wider variety of microbial datasets.

### 3.2 Comparison of phylogenies under different partition sizes

To ensure that downstream results would be similar across different partition sizes, we compared the phylogenies obtained from RAxML [13] using the core variants output by the different Parsnp runs for the bacterial datasets (Table 2). For all three datasets, there were many nearly-identical sequences included which yielded a large number of effectively zero-length branches. To account for this, we used the normalized weighted Robinson-Foulds distance (N-wRF) [14], a branch-length-weighted metric, to compare phylogenies. This is equivalent to the normalized ℒ_1_ distance on the branch lengths, where non-shared branches are treated as having length 0 in the other tree. The wRF distance was normalized by the sum of branch lengths across both trees for each comparison.

**Table 2:** Normalized weighted Robinson Foulds distance between partitioned phylogenies to the non-partitioned phylogeny. Parsnp was executed with different partition sizes *p* on each set of genomes. The resulting phylogenies from each run were compared to the non-partitioned phylogenies using the normalized weighted Robinson-Foulds distance (N-wRF) calculated using DendroPy [10]. ^*∗*^As the non-partitioned version of Parsnp failed to run on the *K. pneumoniae* dataset, we compared the resulting trees to the tree obtained from the *p* = 250.

|  | *M. tuberculosis* | *S. aureus* | *K. pneumoniae* * |
| --- | --- | --- | --- |
| $p = 50$ | 0.094 | 0.120 | 0.068 |
| $p = 100$ | 0.074 | 0.119 | 0.059 |
| $p = 250$ | 0.043 | 0.108 | — |

Except for the *S. aureus* runs, all distances were less than 0.10, signifying high levels of similarity between the phylogenies. In the *S. aureus* runs, the N-wRF distances were only slightly larger than 0.10, likely due to the core-genome alignments being twice as large as the non-partitioned run and slightly more divergent. Altogether, these results show a largely consistent phylogeny across different partition sizes and non-partitioned runs.

### 3.3 Comparison to Parsnp v1

The experiments in Table 1 include a non-partitioned run (i.e. *p* = *n*) which is algorithmically nearly identical to version 1 of Parsnp. Nonetheless, the software has evolved and so a small direct comparison to Parsnp v1 *vis-a-vis* resource usage is presented here.

We obtained the set of 27 *P. tritici-repentis* genomes [15] used to benchmark Parsnp v1 in a recent study [11] and ran Parsnp v2 without partitioning. While most the metrics of Parsnp v2 remain the same as those reported in [11], the memory usage dropped from 57Gb to 8.4Gb.

## 4 Conclusion

Despite approaching the tenth anniversary of its release, Parsnp has an active user-base and remains a leading method for core-genome alignment of large sets of microbial genomes. Parsnp v2 represents the collective improvements from nearly five years of maintenance and development resulting in several new features, bug-fixes, and enhancements to the user experience. In addition, Parsnp v2 includes a new partitioning strategy which leads to substantial reductions in runtime and memory usage. Collectively, the improvements to Parsnp v2 enable alignments of larger datasets in a package that is more efficient, flexible, robust, and convenient than the original version.

## 6 Author contributions statement

B.K. implemented all new additions to Parsnp v2. B.K. and V.H. identified and resolved bugs from Parsnp v1. B.K, E.K, and M.G.N performed the experimental analyses. B.K., M.G.N., A.M.P., and T.J.T. contributed to the manuscript and experimental design. T.J.T supervised the study.

## Acknowledgements

The authors would like to thank Yunxi Liu, Advait Balaji, Yilei Fu, and Ada Ma for their assistance with the development and testing of Parsnp v2.

