## Supplemental Material for "Parsnp 2.0: Scalable Core-Genome Alignment for Massive Microbial Datasets"

January 31, 2024

### 1 Supplementary Materials

#### 1.1 Identifying the final core-genome coordinates from a set of partitioned alignments

In order to identify the coordinates of the final core genome with respect to a reference sequence, we compute the intersection of the aligned reference regions for each partition. Let $q$ be the number of partitions of $n$ query genomes resulting in $p$ genomes per partition, and $R_i = \{(b_1, e_1), (b_2, e_2), ..., (b_m, e_m)\}$ be the begin and end coordinates of the reference for each of the $m$ LCBs in partition $i$.

We compute the interval intersection of all $R_i$ interval sets to obtain the coordinates of the final core-genome intervals, $R^*$. Intervals which are smaller than 10 base pairs are removed from $R^*$. The output alignments of each partition are then trimmed to match the coordinates in $R^*$. At the end of this step, each partition will have an alignment file with $|R^*|$ LCBs, where each LCB corresponds to one of the intervals in $R^*$.

#### 1.2 Merging partitioned alignments

As the merging process is identical across each LCB, we will describe the case for merging a set of LCBs corresponding to the same reference interval. We use two operations, appending and stacking, where appending $LCB_b$ to $LCB_a$ is the process of adding a each column in $LCB_b$ to the end of $LCB_a$, and stacking $LCB_b$ on $LCB_a$ is the process of adding the rows in $LCB_b$ to $LCB_a$. We denote $LCB_i$ as the alignment in partition $i$. Our goal is merge all $LCB_i$ over all $i \in \{1, 2, ..., q\}$.

In the case of LCBs where there are no indels in the reference for every LCB, the task is straightforward: stack all of the LCBs together. However, special care must be taken when indels are present in the reference sequences. Consider column $j$ of $LCB_i$, denoted as $LCB_i[j] = [c_1, c_2, c_3, ...c_{1+p}]$, where $LCB_i[j][k] = c_k$ represents the character in row $k$, column $j$ of $LCB_i$, and row 0 always corresponds to the reference. If

$LCB_i[j][0] =$ " $-$ " and $LCB_i[j][k] \neq$ " $-$ ", then we have no information on how the base at $LCB_i[j][k]$ is related to the query sequences from other partitions.

To remedy this, we assemble the output LCB, $LCB^*$ column by column. At column $j^*$ in $LCB^*$, we keep track of the corresponding column in each of the partition LCBs, denoted by $j_i$, $i \in q$. If $LCB_i[j_i][k] \neq$ " $-$ " and $LCB_i[j_i][0] =$ " $-$ " for any $i \in [q]$, $k \in [p]$, then instead of stacking all columns and then appending to $LCB^*$, we append the stacked columns to a temporary LCB $X$. Once we come across a column where no LCB has any indels in the reference, we compute a multiple sequence alignment $A$ of the sequences in $X$ using SPOA [1], where each sequence is the concatenation of non-gap characters in each row. The alignment $A$ is then appended to $LCB^*$, $X$ is reset to the empty LCB and the process continues.
